# Sleep and gestation duration are correlated, but not with cancer risk or prevalence across vertebrates

**DOI:** 10.64898/2026.06.12.731839

**Authors:** Tanya Mishra, Zachary T. Compton, Stefania E. Kapsetaki

**Author notes:** corresponding author: Email: S.E.K. Emails: T.M., Z.T.C. Primary affiliation of S.E.K.: Hellenic Open University, Patras, Greece.

## Abstract

Cancer prevalence varies across species, with traits such as litter/clutch size, gestation duration, carnivory, and adult mass, partly explaining this variation. Yet no coherent explanation exists for why shorter gestation and carnivory both correlate with cancer prevalence or risk. Given that carnivores sleep more than herbivores, sleep reportedly being a compensation for brain immaturity after gestation, longer sleep appearing in species with shorter gestation, and shorter gestation in species with higher cancer prevalence, we hypothesize that sleep may mediate the gestation-cancer association. We obtained neoplasia prevalence, cancer prevalence, cancer mortality risk (n ≥ 20 individuals/species), gestation duration, and sleep duration data across vertebrates. We tested whether sleep duration mediates the known gestation-cancer prevalence association, whether sleep duration is directly correlated with neoplasia/cancer prevalence or risk, and tested the known gestation-sleep duration association using more vertebrate species and phylogenetic generalized least squares analyses. Gestation and sleep duration were not correlated with neoplasia prevalence, cancer prevalence, or cancer mortality risk. Gestation and sleep duration were negatively correlated. These results highlight the complexity of understanding how multiple physiology variables explain the variation in cancer prevalence or risk across vertebrates, and the need to verify previously known associations with more powerful and robust statistical tools.

## Introduction

There is variation in the prevalence and risk of cancer across species (e.g. [1,2]). Many studies have tried to understand this variation, finding variables that correlate with cancer prevalence or risk across vertebrates. Among these variables, some, more or less consistently, correlate with cancer prevalence or risk. These include litter/clutch size [3–5], gestation duration [2], lactation duration [5], adult body mass [2,6,7], trophic levels/carnivory [1,8], the negative regulation of Transforming Growth Factor beta (TGFβ) production [9], the number of active transposable elements (specifically, long interspersed nuclear element-1 and short interspersed nuclear elements) [10], and germline mutation rates [11]. Apart from life history tradeoffs proposed to explain the positive correlation between litter or clutch size and cancer prevalence or risk [3–5], biomagnification possibly explaining the higher prevalence and risk of cancer in carnivores [1,8], and mutations in cancer-promoting/suppressing genes possibly explaining the positive correlation between germline mutation rates and cancer risk across vertebrates [11], there has not yet been a coherent explanation for why both gestation duration and carnivory correlate with cancer prevalence or risk.

We propose the following series of events to help explain why both gestation duration and carnivory correlate with cancer prevalence or risk. First, prey species having a higher risk of being found and eaten by predators probably evolved sleeping less than predators. As Siegel [12] mentions, giraffes "must not sleep deeply if they are to survive". Second, “immaturity at birth is the single best predictor of REM sleep time throughout life” [13,14], and “the duration of sleep seems to compensate for the incomplete maturity of the brain after gestation” [14]. Third, sleeping less may have evolved in parallel with longer gestation lengths in prey than predators. Fourth, having longer gestation duration may have meant more resources towards somatic maintenance and cancer defences, thus lower cancer prevalence or risk in prey than predators.

Are there any molecular links between these variables? Growth hormone may be one of the key molecules linking the variation in sleep duration, gestation duration, and cancer prevalence or risk across vertebrates. Growth hormone is detected at higher concentrations during sleep [15,16]. This hormone increases cell proliferation, angiogenesis, and metastasis [17–20], and induces secretion of TGFβ in humans and mice [21]. Less negative regulation of TGFβ production appears in species with higher cancer prevalence [9]. Growth hormone also helps in wound healing [22,23], and possibly contributes to wound healing and cancer having common hallmarks [24]. Thus, given that herbivores tend to sleep less than carnivores [25–27], they may be secreting and be exposed to relatively less growth hormone after birth.

Further support of possibly relatively lower secretion/exposure to growth hormone in herbivores comes from reports of slower wound healing in herbivores than carnivores [28], and higher trauma mortality in herbivores than carnivores [28]. A lower exposure to growth hormone in herbivores, versus carnivores, after birth may mean that they are less vulnerable to angiogenesis (a hallmark of cancer [29]) and uncontrollable cell proliferation (also a hallmark of cancer [29]), explaining their relatively lower neoplasia prevalence and cancer risk [1,8].

Correlations further support the majority of these points. Specifically, first, total sleep duration and REM sleep duration is negatively correlated with predation risk (i.e. more exposed environments [30–32]); herbivores are known to sleep less than carnivores (within mammals [25–27]). Second, animals born relatively more mature (precocial animals) than others (altricial animals) have less REM sleep in adulthood [26,33]. Third, mammalian species with shorter sleep duration tend to have longer gestation duration [26,30,32,34,35] (these analyses [30,32,34] were not phylogenetically controlled, whereas Lesku et al.’s analyses were phylogenetically controlled [26,35]). Fourth, vertebrate species with longer gestation length tend to have lower cancer prevalence [2] and incidence/risk of cancer mortality [5]; cancer mortality is known to be lower in non-Carnivora [1]; and neoplasia prevalence is lower in herbivores than primary carnivores when controlling for domestication [8].

Roche et al. highlight the difficulty of untangling the contribution of sleep duration to cancer risk due to sleep duration’s association with many variables [36]. To find further evidence as to whether the above links are true, we sought to fill this gap in the literature, i.e., test whether sleep duration mediates the known association between gestation duration and cancer prevalence/risk, and whether sleep duration is directly correlated with cancer prevalence or risk across vertebrates using Zero-Inflated Beta-Binomial Phylogenetic Generalized Linear Mixed Models (ZiBBPGLMMs). ZiBBPGLMMs take into account the binomial nature of the cancer prevalence and cancer mortality data and are thus more suitable than phylogenetic generalized least squares (PGLS) regression analyses [37] (used in previous studies, e.g. [2–5,8,9,11,38]) for cancer prevalence and cancer mortality risk comparative analyses. We also ran the previously reported association between sleep duration and gestation duration [26,30,32,34,35] to test if it holds using total sleep duration data, more species across vertebrates, and phylogenetically-controlled PGLS. We refer to sleep duration as the total number of hours each species sleeps on average per day.

Based on the above proposed series of associations and evolutionary pressures on the duration of sleep, gestation, and prevalence/risk of cancer in herbivores versus carnivores, we hypothesize that sleep duration mediates the negative correlation between gestation duration and cancer prevalence, sleep duration positively correlates with cancer prevalence or risk, and sleep duration negatively correlates with gestation duration.

## Methods

### Data collection

We obtained aggregated neoplasia and cancer prevalence data [2,3,8,28,39], and cancer mortality risk data [1] for vertebrate species from the literature. The neoplasia and cancer prevalence data [2,3,8,28,39] included mammals, reptiles, amphibians, and birds, whereas the cancer mortality risk data [1] included only mammals.

We obtained average gestation (in the case of non-oviparous animals) or incubation (in the case of oviparous animals) duration (months) data for each species from Compton et al. [2,3] and Animal Diversity Web (https://animaldiversity.org/).

We searched for sleep duration (i.e. total hours of daily sleep) data in the literature for the species for which we had either neoplasia prevalence, cancer prevalence, or cancer mortality risk data. We found sleep duration data of 74 vertebrate species, of which one was a reptile, 15 were birds, and 58 were mammals (Supplementary Material; S2). When there were many reports of sleep duration for a single species in the literature, we obtained the value that was measured from the largest number of individuals in that species. When there were many reports with the same number of individuals, we chose the sleep duration value that was reported for adult individuals. S.E.K. and T.M. validated the sleep duration data (Supplementary Material; S2).

### Statistical analyses

In all analyses we used R version 4.4.2 (2024-10-31) [40] and the caper, phytools, geiger, and tidyverse packages as have been used in previous comparative oncology studies (e.g. [2,8,11,28,39,41]). All analyses were phylogenetically controlled using a phylogenetic tree from TimeTree (Supplementary material; S5). Data on sleep duration, gestation duration, neoplasia prevalence, cancer prevalence, and cancer mortality were not available for every species. Thus, in each analysis we pruned the tree to include only the species that had data for both the dependent and independent variable.

We tested whether sleep duration and gestation duration are correlated with neoplasia prevalence, cancer prevalence, or cancer mortality. In these analyses where the dependent variable is binomial, i.e., neoplasia prevalence, cancer prevalence, or cancer mortality, we performed ZiBBPGLMMs (Supplementary material; S3). A previous study comparing tumor prevalence versus body mass across mammals and birds has also used phylogenetically-controlled beta binomial regressions [42]. The dependent variable in this model takes into account the number of successes (i.e. number of neoplasms, malignancies, or cancer mortality cases depending on the analysis) and the total trials (i.e. the total number of necropsies in the case of the neoplasia and cancer prevalence analyses, or the total number of individuals in the cases of the cancer mortality analyses). We only included species that had neoplasia prevalence, cancer prevalence, or cancer mortality risk data from at least 20 individuals. Such a threshold has also been used in previous studies (e.g. [1–3,8,38]). We performed the ZiBBPGLMM analyses across vertebrates and within separate classes. S.E.K. and T.M. validated the ZiBBPGLMM code and results.

We also tested whether sleep duration and gestation or incubation duration are correlated. Given that sleep duration and gestation duration are not binomial variables, we performed PGLS analyses, instead of ZiBBPGLMM. We used log10-transformed gestation or incubation duration as the independent variable and sleep duration as the dependent variable (Supplementary material; S4). We performed PGLS analyses across vertebrates as well as within separate classes.

## Results

The species used in this study varied in the average duration of gestation or incubation, the total number of hours they slept on average per day, their cancer prevalence, and cancer mortality risk. For example, the common bluetongue *Tiliqua scincoides* has the shortest incubation duration (0.02 months), whereas the African bush elephant *Loxodonta africana* has the longest gestation duration (22 months). The roe deer *Capreolus capreolus* and Gambian pouched rat *Cricetomys gambianus* sleep the least (2.6 hours on average per day), whereas the big hairy armadillo *Chaetophractus villosus* sleeps the most (20.4 hours on average per day). The chaffinch *Fringilla coelebs* is among the species with the lowest cancer prevalence (0%), whereas the Virginia opossum *Didelphis virginiana* has the highest cancer prevalence (51.9%). The Gelada baboon *Theropithecus gelada* is among the species with the lowest cancer mortality risk (0%), whereas the jaguar *Panthera onca* has the highest cancer mortality (24.4%) in the dataset.

### Gestation or incubation duration is not correlated with cancer prevalence or risk

To test whether sleep duration mediates the association between gestation duration and cancer prevalence, we reran the comparison of gestation duration versus cancer/neoplasia prevalence analysis of Compton et al. across vertebrates [2], and gestation duration versus cancer mortality risk of Dujon et al. across mammals [5], though this time using ZiBBPGLMM. We found no correlation between gestation/incubation duration and neoplasia prevalence (Fig. 1A; N = 146; ZiBBPGLMM; slope = –0.05; 95% credible intervals: –0.28, 0.17), or cancer prevalence (Fig. 1B; N = 146; ZiBBPGLMM; slope = –0.09; 95% credible intervals: –0.34, 0.16) across vertebrates. Within mammals, there was no correlation between gestation duration and neoplasia prevalence (Fig. 1A; N = 91; ZiBBPGLMM; slope = –0.21; 95% credible intervals: –0.52, 0.10; Supplementary material; S6), cancer prevalence (Fig. 1B; N = 91; ZiBBPGLMM; slope = –0.23; 95% credible intervals: –0.54, 0.08; Supplementary material; S6), or cancer risk (Fig. 1C; N = 40; ZiBBPGLMM; slope = –0.25; 95% credible intervals: –1.10, 0.57). Within birds, there was no correlation between gestation duration and neoplasia prevalence (Fig. 1A; N = 34; ZiBBPGLMM; slope = 0.25; 95% credible intervals: –0.88, 1.36; Supplementary material; S6) or cancer prevalence (Fig. 1B; N = 34; ZiBBPGLMM; slope = 0.49; 95% credible intervals: –0.85, 1.82; Supplementary material; S6). Within reptiles, there was no correlation between gestation/incubation duration and neoplasia prevalence (Fig. 1A; N = 18; ZiBBPGLMM; slope = –0.10; 95% credible intervals: –0.54, 0.34; Supplementary material; S6) or cancer prevalence (Fig. 1B; N = 18; ZiBBPGLMM; slope = –0.12; 95% credible intervals: –0.56, 0.34; Supplementary material; S6). The absence of a correlation between gestation/incubation duration and neoplasia prevalence, cancer prevalence, or cancer risk remained when controlling for sleep duration (additive model: gestation/incubation duration + sleep duration; Supplementary material; S6).

**Figure 1.**
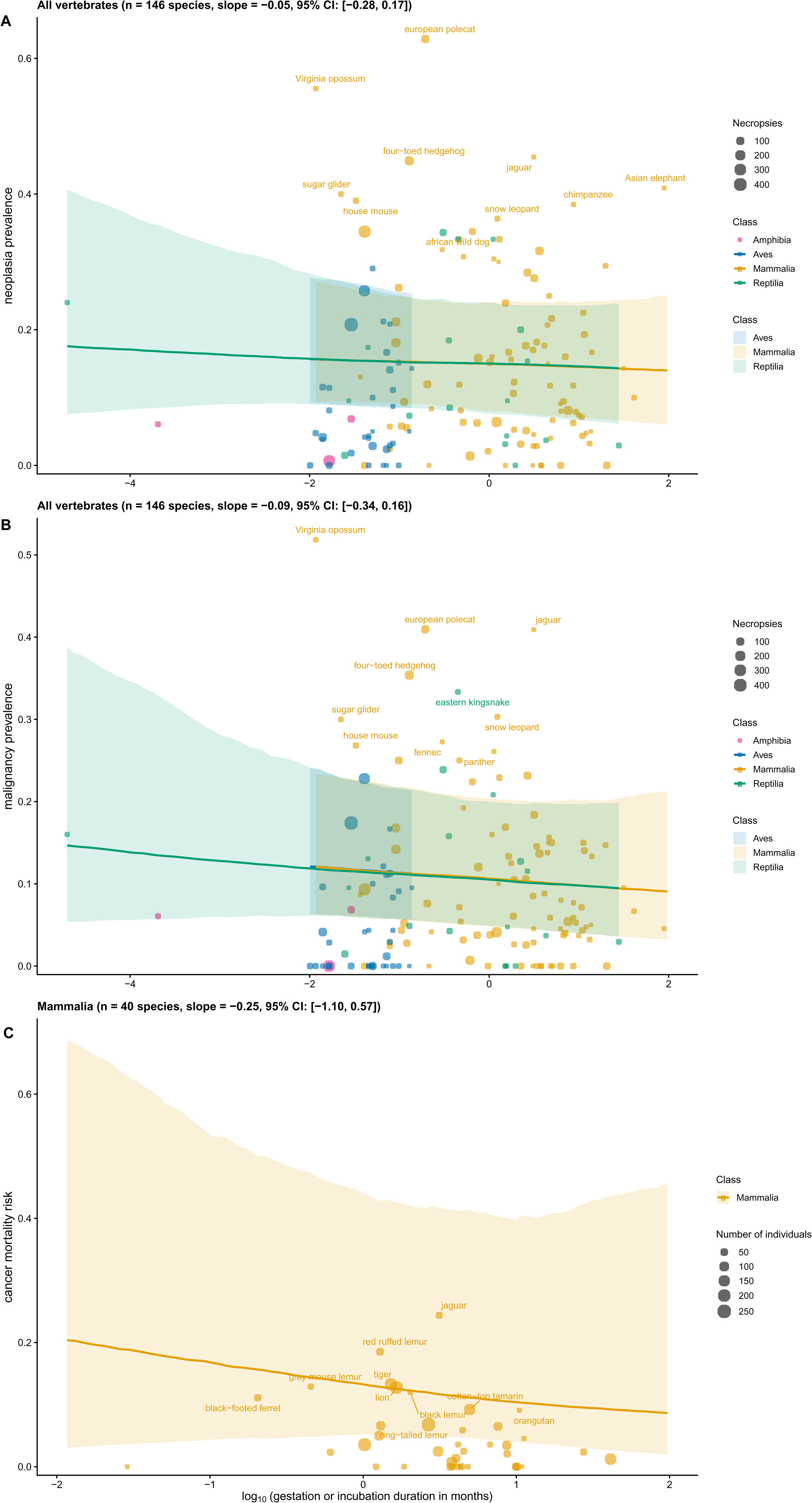
Gestation or incubation duration is not correlated with neoplasia prevalence, cancer prevalence, or cancer mortality risk. The x axis is log10 transformed. Each dot is a different species. Dot size shows the number of necropsies (A, B) or total number of individuals (C) per species. A regression line or ribbon is not shown for Amphibia, since there were only three species of amphibians in these analyses (A, B). We show the common names of the top 10 species with the highest neoplasia prevalence (A), cancer prevalence (B), or cancer mortality risk (C) values. The regression lines are based on the ZiBBPGLMMs. The ribbons represent the 95% credible intervals obtained from the ZiBBPGLMMs. The slope and 95% credible interval values shown in the titles were obtained from the ZiBBPGLMMs. The summary outputs from the statistical analyses of all species and within each class (Aves, Reptilia, Mammalia) are provided in the Supplementary Material (S6).

### Sleep duration is not correlated with cancer prevalence or risk

In order to test whether sleep duration is directly correlated with neoplasia or cancer prevalence or cancer risk, we performed ZiBBPGLMMs of sleep duration versus neoplasia prevalence and cancer prevalence across vertebrates, and cancer risk across mammals. We found that sleep duration is not correlated with neoplasia prevalence (Fig. 2A; N = 38; ZiBBPGLMM; slope = 0.03; 95% credible intervals: –0.07, 0.12) or cancer prevalence (Fig. 2B; N = 38; ZiBBPGLMM; slope = 0.04, 95% credible intervals: –0.06, 0.14) across vertebrates. Within mammals, sleep duration did not correlate with neoplasia prevalence (Fig. 2A; N = 31; ZiBBPGLMM; slope = 0.02; 95% credible intervals: –0.08, 0.13; Supplementary material; S6), cancer prevalence (Fig. 2B; N = 31; ZiBBPGLMM; slope = 0.03; 95% credible intervals: –0.07, 0.13; Supplementary material; S6), or cancer risk (Fig. 2C; N = 28; ZiBBPGLMM; slope = 0.06; 95% credible intervals: –0.07, 0.18). Similarly, within birds, sleep duration did not correlate with neoplasia prevalence (Fig. 2A; N = 6; ZiBBPGLMM; slope = –0.06; 95% credible intervals: –0.54, 0.44; Supplementary material; S6) or cancer prevalence (Fig. 2B; N = 6; ZiBBPGLMM; slope = –0.09; 95% credible intervals: –0.63, 0.49; Supplementary material; S6). We could not perform separate statistical analyses between sleep duration and neoplasia or cancer prevalence for reptiles since there was only one reptile species with both sleep duration and neoplasia or cancer prevalence data.

**Figure 2.**
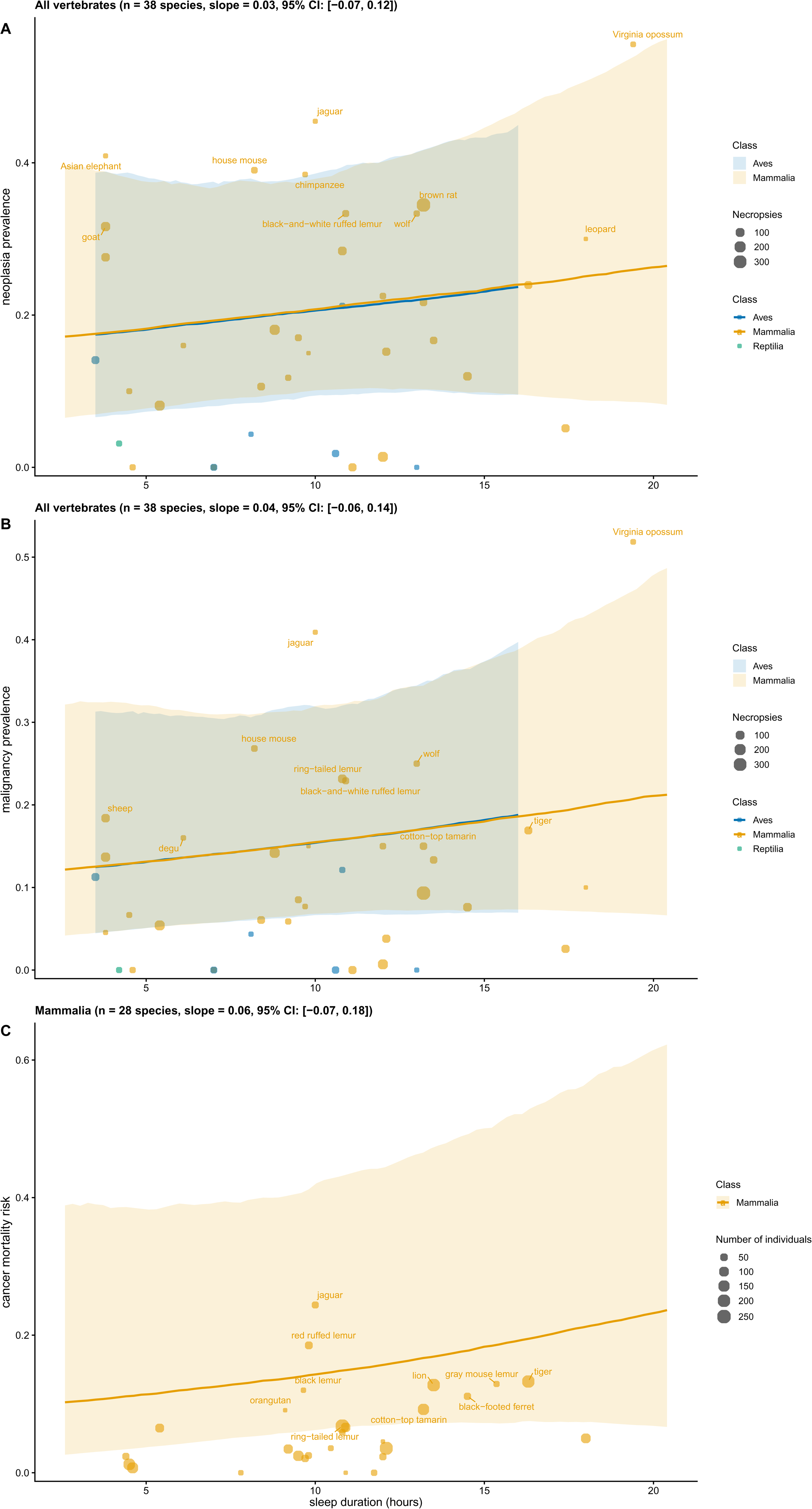
Sleep duration is not correlated with neoplasia prevalence, cancer prevalence, or cancer mortality risk. Each dot is a different species. Dot size shows the number of necropsies (A, B) or total number of individuals (C) per species. A regression line or ribbon is not shown for Reptilia, since there was only one species of reptile in these analyses (A, B). We show the common names of the top 10 species with the highest neoplasia prevalence (A), cancer prevalence (B), or cancer mortality risk (C) values. The regression lines are based on the ZiBBPGLMMs. The ribbons represent the 95% credible intervals obtained from the ZiBBPGLMM. The slope and 95% credible interval values shown in the titles were obtained from the ZiBBPGLMMs. The summary outputs from the statistical analyses of all species and within each class (Aves, Mammalia) are provided in the Supplementary Material (S6).

### Sleep and gestation duration are negatively correlated

To test previous results showing that sleep and gestation duration are negatively correlated [26,30,32,34,35] hold across vertebrates when using PGLS, we performed PGLS analyses between sleep and gestation/incubation duration across vertebrates. We found that sleep and gestation/incubation duration are negatively correlated across vertebrates (Fig. 3; N = 60; PGLS; lambda = 0.89; regression coefficient = –5.21; R² = 0.20; *P*-value = 0.008; 95% confidence intervals: –8.99, –1.44), and in a separate analysis that only included mammals (Fig. 3; N = 50; PGLS; lambda = 0.77; regression coefficient = –5.83; R² = 0.27; *P*-value = 0.003; 95% confidence intervals: –9.52, –2.14), but not within birds (Fig. 3; N = 9; PGLS; lambda = 0.97; regression coefficient = 7.09; R² = 0.07; *P*-value = 0.54; 95% confidence intervals: –19.00, 33.20). We could not perform a separate statistical analysis for reptiles as there was only one species of reptile with both sleep duration and gestation/incubation duration data.

**Figure 3.**
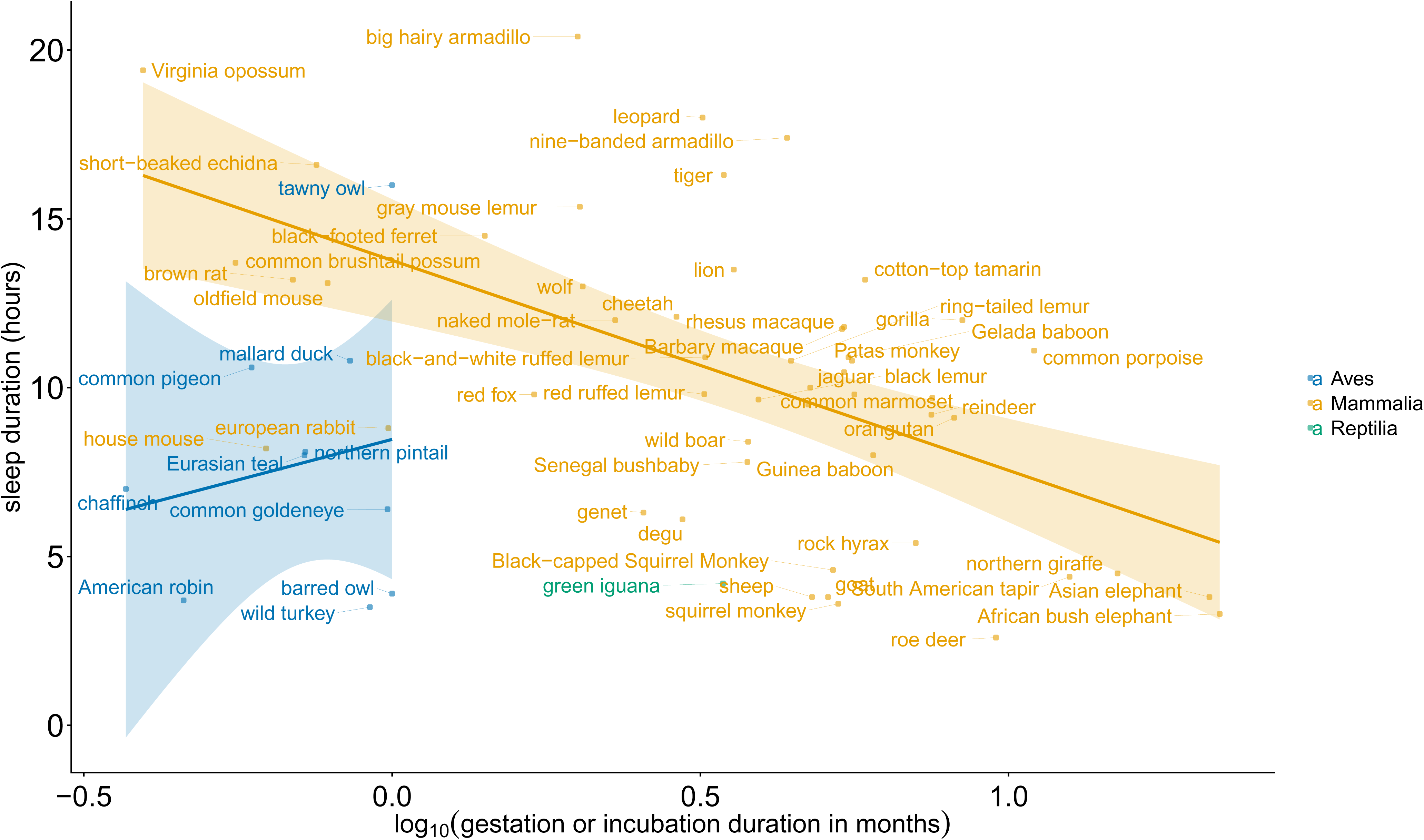
Sleep duration is negatively correlated with gestation or incubation duration. The x axis is log10 transformed. Each dot is a different species. A regression line or ribbon is not shown for Reptilia, since there was only one reptile in this analysis. We show the common names of species that do not highly overlap with neighbouring common names. The regression lines are based on the non-phylogenetically controlled linear model. The ribbons show the 95% confidence intervals of the regression lines.

## Discussion

In this study we aimed to find whether sleep duration mediates the known association between gestation/incubation duration and cancer prevalence [2]/risk [5] across vertebrates, whether there is a direct association between sleep duration and cancer prevalence, and whether sleep duration and gestation/incubation duration are correlated. We found that a relationship between gestation/incubation duration and cancer prevalence across vertebrates does not exist when using ZiBBPGLMMs. There is also no direct association between sleep duration and neoplasia prevalence, cancer prevalence, or cancer risk. Only gestation/incubation and sleep duration were negatively correlated.

Finding or not finding a correlation between gestation/incubation duration and cancer prevalence depends on the statistical analysis. Previous studies using the same neoplasia and cancer prevalence data [2], and cancer risk data [5], but using PGLS, had found a negative association between cancer prevalence/risk and gestation/incubation duration. However, this result did not hold in this study that used ZiBBPGLMM. We found that gestation/incubation duration is not correlated with cancer prevalence, neoplasia prevalence across vertebrates, or cancer mortality risk across mammals. The finding of different statistical significance between these two articles [2,5] versus this study highlights the importance of methodological differences, specifically in the statistical analyses in this case, in comparing the results of different studies. The observation that the negative correlation between gestation or incubation duration and cancer prevalence [2]/risk [5] does not hold in this study, shows that it is not a strong correlation that withstands the use of a different statistical test which takes into account the large number of zero values and binomial nature of the neoplasia prevalence, cancer prevalence, and cancer mortality risk data.

Sleep duration is not correlated with neoplasia prevalence, cancer prevalence, or cancer risk. No previous studies have performed statistical analyses to test whether sleep duration is correlated with neoplasia prevalence, cancer prevalence, or cancer risk across vertebrates. Our finding that there is no such correlation draws our attention to exploring the molecular reasons as to why such an association does not exist, and other variables that might explain the variation in cancer prevalence/risk across vertebrates. For example, litter or clutch size appears to be a variable that relatively more consistently, and among a large number of species, explains part of the variation in cancer prevalence or risk across birds [3] and mammals [4,5].

The finding of a negative correlation between sleep and gestation duration across vertebrates using PGLS is consistent with previous findings in mammals that did not use PGLS. Specifically, Allison and Cicchetti found the same results using fewer species of mammals (39 species) and not controlling for phylogeny [30]. Gonfalone found a similar result using 79 species of mammals, but looking at REM sleep duration and not controlling for phylogeny [34]. Capellini et al. found a similar result when comparing REM sleep duration, non-REM sleep duration, and total sleep duration, versus gestation duration across mammals, though not controlling the analyses for phylogenetic non-independence [32].

Lesku et al. [26,35] found a similar negative correlation between gestation duration and REM sleep duration (54-83 species in their studies; versus 60 species in our study), controlling for phylogeny but using a slightly different package than we used. The COMPARE 4.6b [43] programme that they used also uses Brownian motion approaches, but does not enable the measurement of phylogenetic signals. The observation that gestation duration and sleep duration remain negatively correlated across multiple studies, indicates that so far it is a robust correlation that persists when using a variety of different statistical methods.

### Limitations & Future directions

In some analyses (Fig.s 2, 3: birds and reptiles) there are currently too few species to draw robust conclusions. Sleep duration showed a positive trend with neoplasia prevalence, cancer prevalence and cancer risk, and gestation/incubation duration showed a negative trend with neoplasia prevalence, cancer prevalence, and cancer risk, though these analyses were not statistically significant. Future studies should investigate the duration of sleep across many thousands more species and across many more classes, as highlighted also in Siegel’s review on sleep studies across invertebrates and vertebrates [44], in order to test whether gestation/incubation duration and cancer prevalence/risk are correlated, sleep duration mediates associations between gestation/incubation duration and cancer prevalence/risk, and whether sleep duration is directly correlated with cancer prevalence/risk.

The exact duration of sleep of the same animals for which we have cancer data is unknown. We also do not know the exact living conditions of the animals for which we have necropsy data. It would be useful to have these data to be able to control our analyses for high environmental temperature and presence of ectoparasites, both of which are known to reduce sleep duration, in king penguins [45] and great tits [46], respectively. Future studies should track environmental conditions, gestation/incubation duration, sleep duration, and cancer pathology data from the exact same animals, so that a comparison between these variables can be performed in more controlled settings. This is a common limitation in hitherto large-scale comparative oncology studies (e.g. [1,2,5,6,8]) where the physiology variables examined, such as maximum lifespan, average adult weight, and gestation/incubation duration are average species values, and not the average values from the exact same animals from which the cancer data have been obtained.

The correlation between gestation duration and sleep duration does not mean causation. Examining the molecular pathways underlying this correlation would provide further insights as to why this correlation exists. As we mentioned in the introduction, growth hormone may be key in this association due to its concentration often increasing during sleep [15,16] and its function contributing, among others, to cell proliferation [17–20].

Future studies should also test whether correlations between gestation/incubation duration, sleep duration, and cancer prevalence/risk exist when examining species from the wild, and whether there are differences in the associations between these variables in wild versus captive species. For example, several species, such as sloths [47,48] and elephants [49–51], are known to sleep less in the wild than in captivity.

## Conclusions

Our results highlight that the proposed series of events, mentioned in the introduction, regarding explaining the variation in cancer prevalence/risk based on trophic levels, sleep duration, and gestation/incubation duration is not straightforward. Even though gestation/incubation duration and sleep duration are negatively correlated, they are not correlated with neoplasia prevalence, cancer prevalence, or cancer mortality risk. Each variable does not explain 100% of the variation in another variable. For example, gestation/incubation duration explains only 20% (R² = 0.20) of the variation in sleep duration across vertebrates. There is room for future studies to expand the number of species and test new ecophysiology variables that will help explain the variation in cancer prevalence and risk across vertebrates.

## Data & code availability

We provide all data and code used in this study in the supplementary material. The supplementary material will be provided in the published version of the article.

## Statements & Declarations

## Acknowledgements

We thank Vahid Nikoonejad Fard for the original version of the ZiBBPGLMM code which S.E.K. then adjusted for running the specific variables of this project.

## Author contributions

T.M. collected most of the sleep duration data and validated the analyses. S.E.K. conceived the idea for this project, validated the sleep duration data, collected a few more, ran the analyses, made the figures, and wrote the first draft of the manuscript. All authors edited the manuscript.

